# Selective encoding failure of self-face identity in subthreshold depression

**DOI:** 10.64898/2026.05.04.721614

**Authors:** Mao Wen, Binyuan Su, Yufan Chen, Tianyi Gu, Pengmin Qin

**Author notes:** Corresponding author: Pengmin Qin. Mao Wen and Binyuan Su contributed equally to this work and should be considered co-first authors.

## Abstract

Subthreshold depression is associated with significant functional impairment and elevated risk of major depressive disorder. A negative self-concept may disrupt the implicit positive association evoked by one’s own face, impairing incidental encoding of self-relevant information. Whether subthreshold depression involves a selective deficit in encoding self-face identity remains unclear. The attribute amnesia paradigm is well suited to address this question because it can dissociate attentional selection from working memory encoding. Using this paradigm, we examined the issue across two experiments. Experiment 1 employed nonsocial stimuli (animal drawings) and confirmed an intact attribute amnesia effect in subthreshold depression (*n* = 30) comparable to healthy controls (*n* = 30), ruling out a generalized encoding deficit. Experiment 2 replaced targets with faces (self or other) and revealed a selective enhancement of the attribute amnesia effect for self-face identity in subthreshold depression. Specifically, on the surprise trial, accuracy for self-face identity dropped to near-chance levels in the subthreshold depression group, whereas no such deficit emerged for other-faces or in controls. Encoding recovered rapidly once explicit memory expectations were introduced, indicating intact basic encoding capacity. These findings suggest that subthreshold depression is associated with a specific impairment in incidentally encoding self-face identity. This impairment likely stems from a negative self-concept that weakens self-face salience under incidental encoding conditions. By capturing this selective encoding failure, the present study reveals that the self-processing deficit in subthreshold depression can arise at the gating stage between attention and working memory consolidation.

## Introduction

Subthreshold depression refers to a condition characterized by clinically relevant depressive symptoms that do not meet the diagnostic criteria for major depressive disorder (MDD) (Cuijpers & Smit, 2004). Most studies now view depression as a continuum, with subthreshold depression and MDD reflecting the same biopsychological process differing only in severity (Angst et al., 2000; Cuijpers et al., 2013). From a clinical perspective, subthreshold depression is not benign: it is associated with significant functional impairment, an elevated risk of progressing to full-syndrome MDD, and increased rates of suicidal ideation and behavior (Wesselhoeft et al., 2013; Cuijpers & Smit, 2004; Angst et al., 2000).

Importantly, individuals with subthreshold depression may exhibit biases or deficits in self-relevant processing (e.g., self-face processing), a pattern that has been well-documented in clinically depressed populations (Quevedo et al., 2018). Self-processing involves the perception and memory of oneself (Liu et al., 2022) and is modulated by one’s self-concept (Morin, 2006). Self-face processing is considered a fundamental modality of self-processing (Bortolon & Raffard, 2018), and self-face recognition is typically associated with positive self-perception (Blackwood et al., 2003). According to the implicit positive association theory (Ma & Han, 2010), viewing one’s own face automatically activates implicit positive self-attributes, thereby facilitating behavioral responses to the self-face. However, a hallmark of depression is a negative self-concept (Beck, 1976). As a core evaluative component of self-concept, low self-esteem, defined as the lack of positive self-evaluation, extends across the full spectrum from subthreshold depression to major depressive disorder (Hilbert et al., 2019; Zheng et al., 2014) and serves as a causal risk factor for depression-related cognitive abnormalities (Orth & Robins, 2013). Low self-esteem may disrupt the implicit linkage between the self-face and positive self-perception, thereby leading to anomalies in self-face processing among individuals with subthreshold depression. Consistent with this reasoning, depressed individuals show greater avoidance of positive facial stimuli (Gotlib et al., 2004; Hankin et al., 2010), faster responses to negative self-relevant stimuli (e.g., a sad self-face; Fritzsche et al., 2010), and the influence of negative self-concept can extend to the processing of one’s own face (Ma & Han, 2010). Moreover, depression is associated not only with negative self-perception but also with a tendency to view others more positively (Kuiper et al., 1982), suggesting a specific deficit in self-face processing.

Early attentional and perceptual processing of the self-face appears to remain largely intact in subthreshold depression. For instance, depressive traits do not modulate the self-face advantage in visual search tasks, with individuals still searching for their own face faster and more accurately than for other faces (Lee et al., 2024). Moreover, in tasks requiring explicit judgments of self-similarity, individuals with subthreshold depression show recognition accuracy comparable to that of healthy controls (Quevedo et al., 2018). These findings indicate that the observed anomalies in self-face processing are unlikely to stem from impaired attention or perception. Instead, such anomalies may originate from deficits in selective encoding or gating mechanisms within working memory. In subthreshold depression, low self-esteem and a negative self-concept may weaken the implicit positive association typically evoked by one’s own face (Ma & Han, 2010), rendering its identity difficult to gate into working memory under incidental encoding conditions. Consequently, the deficit is more likely to reside at the gating stage between attention and consolidation. Taken together, previous research suggests that subthreshold depression may be associated with impaired selective encoding of self-relevant information in working memory, that is, a tendency toward “insufficient encoding of self-face identity.”

Existing studies on self-face processing in subthreshold depression have largely focused on emotional face perception or explicit recognition tasks (Quevedo et al., 2018). Although the self-face advantage in visual search appears to be preserved (Lee et al., 2024), such tasks cannot dissociate attentional selection from subsequent working memory encoding (cf. Firestone & Scholl, 2015). Therefore, to isolate the incidental encoding of self-face identity, we employed the attribute amnesia paradigm (Chen & Wyble, 2015a). Attribute amnesia (AA) is a counterintuitive phenomenon recently identified in the domain of working memory and conscious awareness. In a typical AA paradigm, participants perform a primary task that requires them to rely on a specific, task-defining attribute of a target stimulus (e.g., its category or a salient feature) to complete each trial—for example, locating a target letter among three distractor numbers. After a number of such “pre-surprise” trials, on a single “surprise” trial participants are unexpectedly asked to report another attribute of the same target (e.g., its identity or color) before they carry out the usual response. Even though participants must have consciously perceived and used that attribute during the task, their accuracy on this unexpected memory test is strikingly low. Importantly, once they become aware that such a report may be required, their performance on the same attribute in the immediately following control trials improves dramatically (Chen & Wyble, 2015a, 2015b, 2016; Chen et al., 2019). This phenomenon, termed attribute amnesia, reveals that mere attention to and awareness of a stimulus attribute do not guarantee its subsequent reportability; instead, the encoding of that attribute into working memory is critically dependent on the observer’s expectation of having to remember it. Recent multimodal evidence suggests that this dissociation is achieved through active inhibitory control mechanisms that gate the transition of attended representations into working memory (Liu et al., 2025). By employing this paradigm, we were thus able to directly examine whether subthreshold depression selectively impairs the incidental encoding of self-face identity into working memory.

We conducted two experiments. Experiment 1 employed nonsocial stimuli (black-and-white line drawings of animals) to establish the baseline robustness of the attribute amnesia effect in individuals with subthreshold depression and to rule out potential confounds arising from general cognitive impairment. Experiment 2 extended this investigation by introducing self-face and other-face stimuli as critical targets, thereby directly examining whether subthreshold depression is associated with a specific deficit in encoding self-face identity into working memory. By comparing performance on the surprise trial with that on the immediately following control trials across the two groups, we sought to elucidate the unique impact of subthreshold depressive traits on the selective encoding of self-relevant information.

### Experiment 1: No generalized encoding deficit for nonsocial stimuli

Experiment 1 was designed to establish the baseline comparability of the attribute amnesia (AA) effect between individuals with subthreshold depression and healthy controls when processing emotionally neutral, nonsocial stimuli. Adopting the standard AA design used in most prior investigations (Chen & Wyble, 2015a), each participant completed a sequence of 160 trials comprising 155 pre-surprise trials, one critical surprise trial, and four subsequent control trials. We examined memory for animal line drawings, which served as a nonsocial control condition. The absence of any group difference in AA magnitude for these stimuli would indicate that subthreshold depression does not entail a generalized deficit in selective encoding.

## Method

### Sample size

We conducted an a priori power analysis using G*Power 3.1 (Faul et al., 2007) to determine the required sample size for all experiments reported in this study. The effect size for the attribute amnesia effect (φ) was estimated at 0.50 based on findings from Chen and Wyble (2015). This power calculation indicated that a minimum of 20 participants was required to detect such an effect with 80% statistical power (α set to 0.05). We set the sample size at 30 participants for each group in this study. No participant was permitted to complete more than one experiment in the study.

### Participants

All participants completed the Beck Depression Inventory-I (BDI-I) and the Chinese Shortened Version of the Autism-Spectrum Quotient (15-item AQ-CSV) prior to the formal experiment. Participant selection criteria followed those used in Mahmoud (2023) and Xu et al. (2025). Participants with 11 ≤ BDI-I ≤ 16 (inclusive) and an AQ-CSV score < 39 were included in the subthreshold depression group, while those with a BDI-I score < 11 and an AQ-CSV score < 39 were included in the healthy control group. A total of 60 university students were recruited, with 30 participants in each group, all aged 18 to 25 years. All participants had normal or corrected-to-normal visual acuity, were right-handed, and had no prior experience with similar experiments. This study was approved by the local review board for the ethical treatment of human participants.

### Apparatus and Stimuli

Stimuli were presented on a 14-inch Lenovo monitor with a resolution of 2880 × 1800 pixels. The experiment was programmed using MATLAB (The MathWorks, Natick, MA) with the Psychophysics Toolbox extensions (Brainard, 1997; Pelli, 1997). Participants sat 50 cm from the screen with their heads stabilized by a chin rest and made responses via a computer keyboard.

All visual stimuli used in this experiment were selected from the standardized line drawing database developed by Snodgrass and Vanderwart (1980). A total of 16 black-and-white line drawings were selected, consisting of 12 object drawings and 4 animal drawings. The object drawings included a shoe, a sofa, a table, corn, a hat, a flower, a bicycle, an eye, a foot, a light bulb, and grapes. The animal drawings included a cat, a rabbit, a rooster, and a dog. All 16 images were presented to each participant. Each stimulus array contained one target (a black-and-white line drawings of an animal) and three distractors (black-and-white line drawings of objects). The experimental materials were randomly assigned across trials. All drawings were presented at a size of 300 × 300 pixels.

### Design and paradigm

As shown in Figure 1, each trial began with a fixation display consisting of a black central fixation cross (0.62° of visual angle) and four black placeholder circles (0.62° × 0.62°) located at the four corners of an invisible square (6.25° × 6.25°). The duration of the fixation display varied between 800 ms and 1800 ms. . After that, the stimulus array appeared for 1000 ms, which was replaced by the masks for 100 ms. Following a 500-ms central black fixation cross that appeared after the mask, four black digits (1–4) appeared at the locations of the four placeholders. Participants were required to judge the location of the target animal picture by pressing the corresponding numeric keys (“1”, “2”, “3”, or “4”). These digits remained on the screen until the participant responded. After the response, a 500-ms correctness feedback was presented. In this experiment, each participant completed a total of 160 trials. The first 155 trials followed the standard procedure (as described above). In the 156th trial (i.e., surprise trial), participants were unexpectedly required to select the target animal picture from four animal pictures presented simultaneously on the screen. Digits 5–8 appeared to the right of the four pictures, and participants pressed the corresponding key (5–8) to indicate the target. After this response, they completed a regular location judgment task as in previous trials. Following the surprise trial, the participants received four more control trials in the same format as the surprise trial.

**Figure 1.**
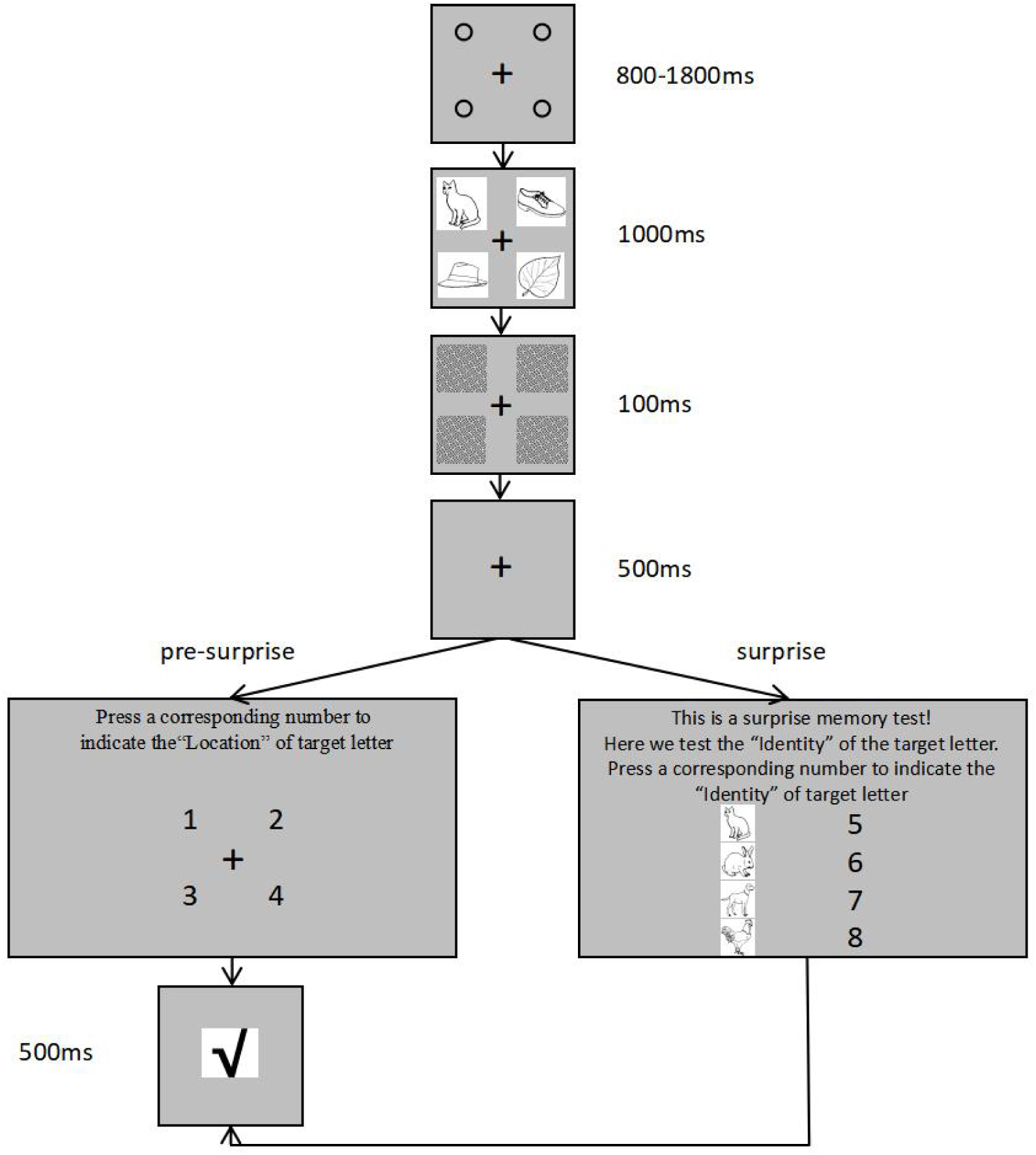
Schematic illustration of the trial sequence in Experiment 1. In pre-surprise trials, participants reported the location of the target animal picture among three distractor object drawings. In the surprise trial, participants were unexpectedly required to report the identity of the target animal picture before making the location judgment. Control trials followed the same format as the surprise trial. The true scale of the stimuli is smaller than depicted; size is for illustration purposes only.

## Results

The results of this experiment are depicted in Table 1. The accuracy of the location report task in pre-surprise trials was 90.00% for the subthreshold depression group and 100.00% for the healthy control group, indicating that all participants could easily select the target animal drawing among distractor object drawings and report its location with a high accuracy. The attribute amnesia effect was observed in both the subthreshold depression group and the healthy control group. For the subthreshold depression group, 15 of 30 (50.00%) participants correctly reported the identity of the animal line drawings in the surprise test, and their performance was significantly lower in the first control trial(93.33%), 50.00% versus 93.33%, χ² (1, *N* = 60) = 11.00, *p* = 0.001, *φ* = 0.61. Similarly, for the healthy control group, the identity accuracy in the surprise test (13 of 30; 43.33%) was also significantly worse than the first control trial (86.67%), 43.33% versus 86.67%, χ² (1, *N* = 60) = 13.00, *p* = 0.000, *φ* = 0.66. The accuracy of the identity task across the remaining three control trials (i.e., the second, third, and fourth) was consistently high: for the subthreshold depression group, 96.67%, 100.00%, and 100.00% correct; for the healthy control group, 96.67%, 100.00%, and 100.00% correct. These results indicate that the poor accuracy on the identity report task was not caused by the surprise trial itself, nor by the failure of conscious perception, but by the failure to selectively encode the information into working memory(Wang et al., 2021).

**Table 1.** Accuracy in Experiment 1 (30 participants in each group)

| Task | Group | Pre-surprise | Surprise | Control 1 | Control 2 | Control 3 | Control 4 |
| --- | --- | --- | --- | --- | --- | --- | --- |
| Location | Healthy control | 100.00% | 73.33% | 90.00% | 96.67% | 96.67% | 96.67% |
|  | Subthreshold depression | 90.00% | 70.00% | 83.33% | 93.33% | 96.67% | 93.33% |
| Identity | Healthy control | N/A | 43.33% | 86.67% | 96.67% | 100.00% | 100.00% |
|  | Subthreshold depression | N/A | 50.00% | 93.33% | 96.67% | 100.00% | 100.00% |

Note. N/A = not available. Controls 1-4 correspond to the four control trials.

### Experiment 2: Selective encoding failure of self-face identity in subthreshold depression

Experiment 1 demonstrated that individuals with subthreshold depression exhibit an intact attribute amnesia effect for nonsocial stimuli, ruling out a generalized encoding deficit. Experiment 2 examined whether subthreshold depression is instead associated with a specific impairment in encoding self-relevant information. Adopting the identical AA design used in Experiment 1 (Chen & Wyble, 2015a), we replaced the animal targets with face stimuli (self-face or other-face) and directly compared the magnitude of the attribute amnesia effect between the two face types across groups. This allowed us to isolate whether the selective encoding deficit in subthreshold depression is uniquely tied to the self-relevance of the to-be-encoded information.

### Method

Experiment 2 was identical to Experiment 1, with the following exceptions. Another 120 participants completed this experiment. Participants were assigned to either the self-face group or the other-face group based on whether they saw a self-face or an other-face as the target in the surprise trial. Specifically, the subthreshold depression group consisted of 30 participants in the self-face condition and 30 in the other-face condition; the healthy control group also included 30 participants in the self-face condition and 30 in the other-face condition. The visual stimuli comprised the same 12 black-and-white line drawings of objects as in Experiment 1, along with four face images: three unfamiliar faces of the same gender as the participant and one self-face. All images were cropped to a circular shape, resized to 300 × 300 pixels, and converted to grayscale. We used the SHINE toolbox (Willenbockel et al., 2010) to adjust the luminance and contrast of these images to control for low-level physical properties (see Figure 2 for example stimuli). In the first 155 trials, participants were asked to report the location of the target face among three distractor object line drawings. Then, in the surprise trial, participants were unexpectedly required to report the identity of the target face before reporting its location. Finally, as before, four more control trials were given after the surprise trial.

**Figure 2.**
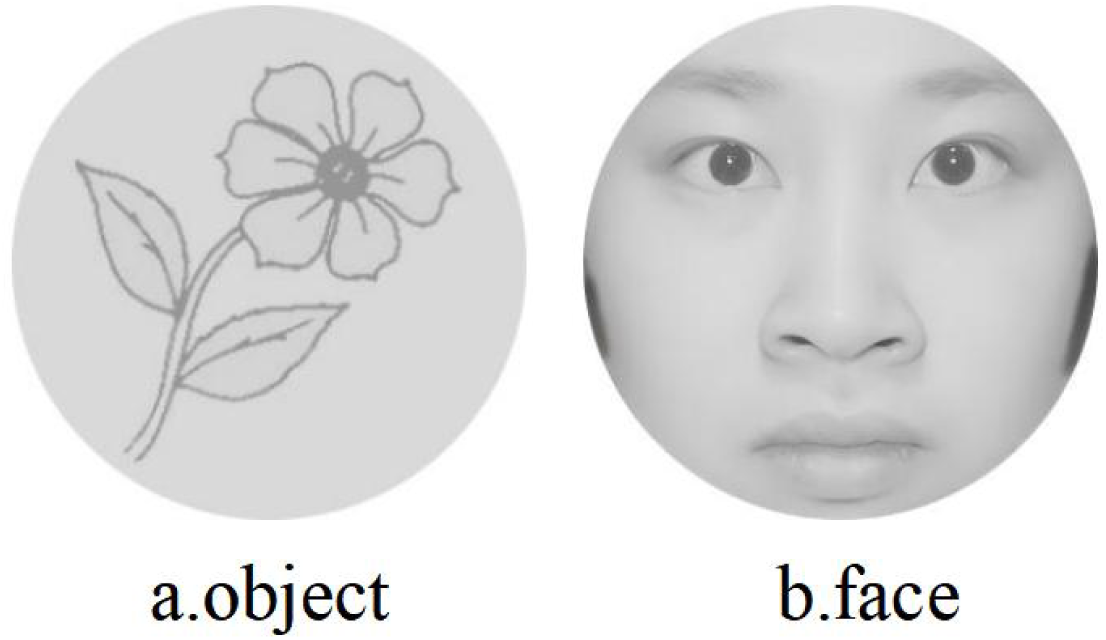
Example stimuli used in Experiment 2. (a) Object line drawings. (b) Face stimuli. The face image is that of one of the authors.

## Results

As shown in Table 2 and Figure 3 and 4, both the subthreshold depression group and the healthy control group exhibited a robust attribute amnesia effect, as evidenced by significantly lower accuracy in the surprise trial than in the second control trial (all *p* < 0.022). For the subthreshold depression group, the accuracy for self-faces (9/30, 30.00%) was significantly lower than that for other-faces (18/30, 60.00%), χ² (1, *N* = 60) = 5.455, *p* = 0.020, *φ* = 0.30. In contrast, for the healthy control group, no significant difference was found between self-faces (15/30, 50.00%) and other-faces (19/30, 63.33%), χ² (1, *N* = 60) = 1.086, *p* = 0.297, *φ* = 0.135. Importantly, identity accuracy on the remaining three control trials (i.e., the second, third, and fourth) was consistently high across all conditions, reaching near-ceiling levels and confirming that the attribute amnesia effect was confined to the surprise trial.

**Figure 3.**
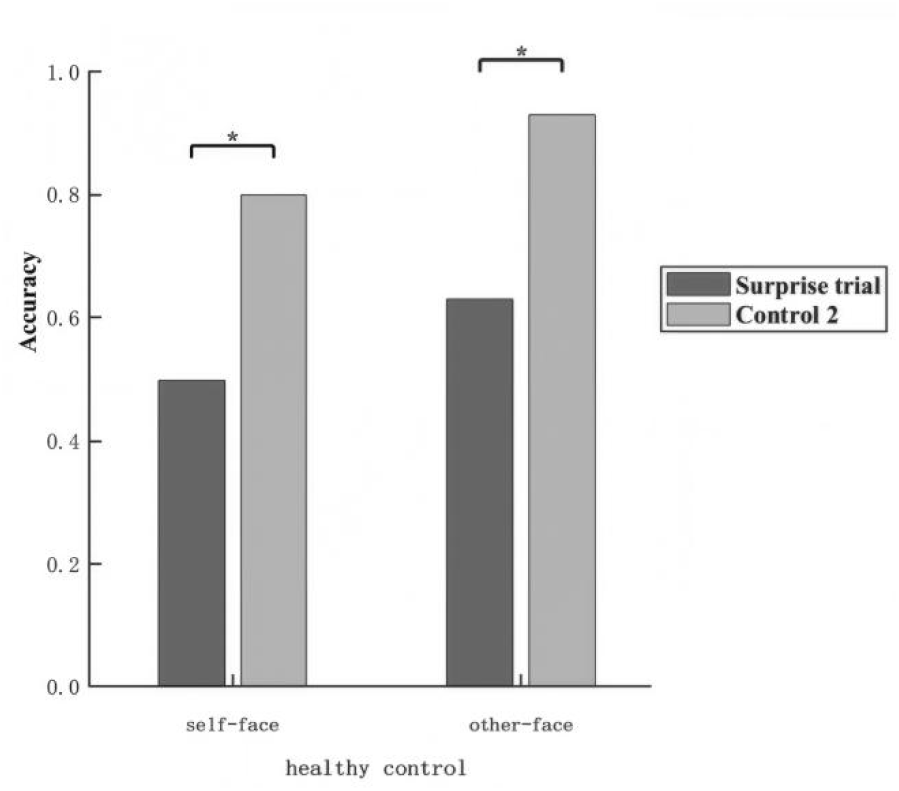
Identity report accuracy in Experiment 2 as a function of face type and trial type for the healthy control group. Accuracy on the surprise trial and the second control trial (Control 2) are shown separately for self-face and other-face conditions. *p < .05. **p < .01. ***p < .001. Control 2 = the second control trial.

**Figure 4.**
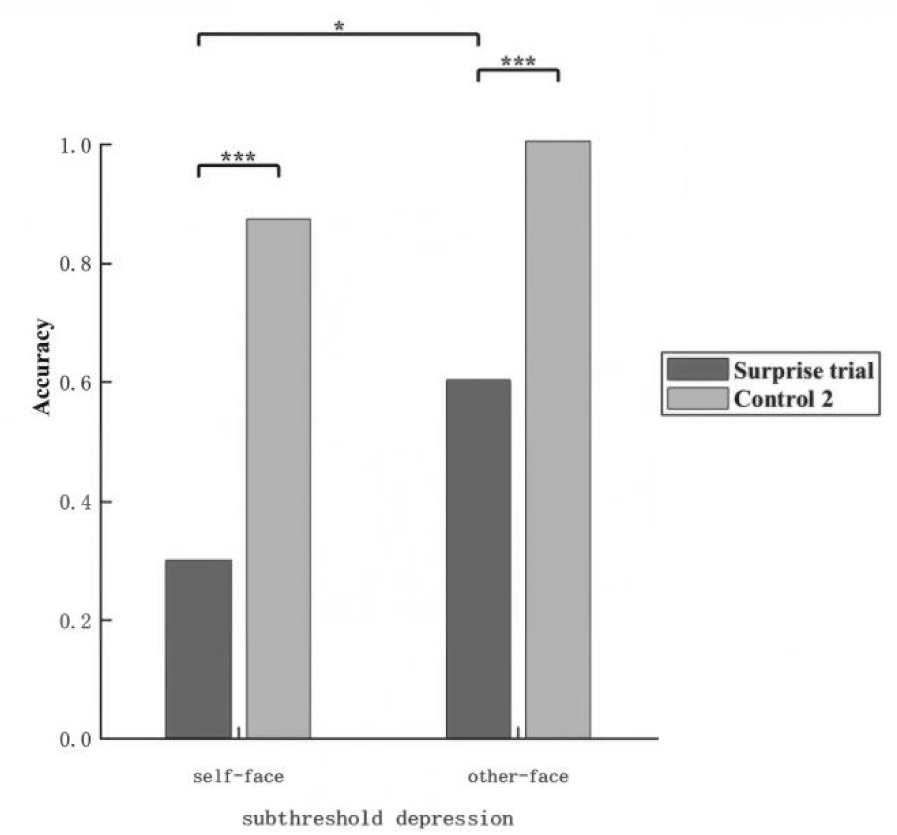
Identity report accuracy in Experiment 2 as a function of face type and trial type for the subthreshold depression group. Accuracy on the surprise trial and the second control trial (Control 2) are shown separately for self-face and other-face conditions. *p < .05. **p < .01. ***p < .001. Control 2 = the second control trial.

**Table 2.** Accuracy in Experiment 1 (30 participants in each group)

| Task | Group | Face<br>type | Pre-surprise | Surprise | Control<br>1 | Control<br>2 | Control<br>3 | Control<br>4 |
| --- | --- | --- | --- | --- | --- | --- | --- | --- |
| Location | Healthy<br>control | self | 100.00% | 66.67% | 90.00% | 96.67% | 100.00% | 93.33% |
|  |  | other | 96.67% | 60.00% | 93.33% | 100.00% | 96.67% | 90.00% |
|  | Subthreshold<br>depression | self | 96.67% | 50.00% | 90.00% | 96.67% | 96.67% | 86.67% |
|  |  | other | 100.00% | 50.00% | 90.00% | 100.00% | 100.00% | 100.00% |
| Identity | Healthy<br>control | self | N/A | 50.00% | 80.00% | 80.00% | 90.00% | 86.67% |
|  |  | other | N/A | 63.33% | 56.67% | 93.33% | 96.67% | 86.67% |
|  | Subthreshold<br>depression | self | N/A | 30.00% | 86.67% | 100.00% | 96.67% | 90.00% |
|  |  | other | N/A | 60.00% | 50.00% | 100.00% | 96.67% | 100.00% |

Note. N/A = not available. Controls 1-4 correspond to the four control trials.

## General discussion

In this study, to examine whether subthreshold depression is associated with a selective deficit in incidentally encoding self-face identity, we employed the attribute amnesia paradigm and compared the accuracy for self-faces with that for other-faces and nonsocial stimuli in the surprise test. Experiment 1 adopted the standard AA design of 155 pre-surprise trials and confirmed that individuals with subthreshold depression exhibited an intact AA effect for animal drawings, comparable to that of healthy controls. In Experiment 2, when the targets were faces, the subthreshold depression group showed a substantially enhanced AA effect specifically for self-faces, with accuracy dropping to near-chance levels, whereas no such difference emerged for other-faces or in the healthy control group, indicating that this selective encoding deficit is uniquely tied to the self-relevance of the information.

### The selective encoding failure for self-face identity in subthreshold depression

The present findings reveal that individuals with subthreshold depression showed significantly lower identity report accuracy for their own face on the surprise trial compared with other-face trials, dropping to near-chance levels, whereas healthy controls exhibited no such difference. Concurrently, Experiment 1 demonstrated no group difference in the magnitude of the attribute amnesia effect for nonsocial animal line drawings.

This pattern of results indicates a selective encoding failure specific to self-face processing in subthreshold depression. Prior behavioral studies on self-face processing in subthreshold depression have focused primarily on early attentional orienting or explicit recognition. Specifically, Lee et al. (2024) found that the self-face advantage in visual search tasks was not modulated by depressive traits, and the self-face advantage is known to occur at an early attentional stage (Jublie & Kumar, 2021). Quevedo et al. (2018) further demonstrated that depressed adolescents performed comparably to healthy controls in both recognition accuracy and reaction time when making explicit judgments of facial self-similarity. Together, these findings indicate that early attentional orienting toward, and explicit recognition of, the self-face remain largely intact in subthreshold depression. Critically, however, these prior studies did not examine the stage at which self-face identity information is gated into working memory. By employing the attribute amnesia paradigm, the present study was able to isolate this specific stage. Our results reveal that, despite intact attention and explicit recognition, individuals with subthreshold depression still fail to encode self-face identity under incidental encoding conditions. This finding localizes the self-processing deficit in subthreshold depression to the gating stage between attention and working memory consolidation, and demonstrates that this deficit persists even for neutral self-faces in the absence of explicit emotional demands.

Notably, neuroimaging evidence from Quevedo et al. (2016, 2018) reveals a critical dissociation: despite normal behavioral performance, depressed adolescents exhibit reduced activation in the medial prefrontal cortex (BA10) and limbic regions (bilateral amygdala, hippocampus) when passively viewing their own happy face, and this aberrant response pattern persists after controlling for depression severity. These regions are centrally involved in detecting emotional salience and consolidating memory; their hypoactivation suggests that the intrinsic salience signal of the self-face is already diminished at the neural level in subthreshold depression. Even though compensatory mechanisms may maintain performance during explicit tasks, this weakened salience signal is insufficient to trigger gating into working memory under incidental encoding conditions.

Connectivity evidence provides further support for functional abnormalities in the gating circuitry. Oh et al. (2025) reported that, during self-face relative to other-face processing, depressed adolescents exhibited heightened bilateral amygdala connectivity with the anterior cingulate cortex, superior temporal gyrus, and frontal gyri compared with healthy controls, a pattern that was attenuated following neurofeedback training. The authors interpreted this hyperconnectivity as reflecting a preponderance of limbic emotional input at the expense of relatively deficient prefrontal executive control. Quevedo et al. (2020) found that during explicit up-regulation of self-processing, depressed adolescents preferentially engaged right amygdala-ipsilateral prefrontal circuits, compensatorily relying on implicit emotion regulation pathways, whereas healthy adolescents relied more on left-lateralized explicit pathways. Collectively, these findings indicate that even under task-engaged conditions, the amygdala-cortical gating circuit in depression is characterized by aberrant lateralization and hyper-coupling. Alarcón et al. (2019) further demonstrated that neural anomalies in self-face processing among depressed adolescents do not vary as a function of facial emotion, as evidenced by enhanced amygdala connectivity with face-processing regions across happy, sad, and neutral faces, suggesting that viewing one’s own face may demand heightened sensitivity or greater cognitive effort.

Consistent with this interpretation, Quevedo et al. (2019) found that during active recall of positive autobiographical memories coupled with neurofeedback, depressed adolescents showed greater activation in the fusiform gyrus, cuneus, and inferior parietal lobule than healthy controls, directly indicating that depressed individuals need to mobilize additional cognitive resources when actively processing the self-face. Liu et al. (2023) further demonstrated, using a modified Sternberg task, that individuals with subthreshold depression exhibit enhanced mid-frontal alpha power during the retention period. Given that alpha power is thought to reflect inhibitory processes that suppress task-irrelevant information (Bonnefond & Jensen, 2012; Jensen et al., 2002; Segrave et al., 2010) and that such inhibition is central to selective encoding into working memory (Miller & Cohen, 2001), this finding points to an abnormality in selective encoding among individuals with subthreshold depression. Moreover, meta-analytic evidence reveals aberrant activation in key regions of the basal ganglia–prefrontal circuit in subthreshold depression, including increased activation in the right lenticular nucleus, putamen, and striatum, and decreased activation in the left lenticular nucleus and putamen (Lyu et al., 2024). The basal ganglia–prefrontal circuit constitutes the canonical neural substrate for selectively gating information into working memory (O’Reilly & Frank, 2006), and the aforementioned regions are integral components of this gating circuitry.

In conclusion, the incidental encoding failure of self-face identity in subthreshold depression likely stems from two interrelated factors. First, negative self-concept and low self-esteem weaken the intrinsic salience signal of one’s own face (Beck, 1976; Ma & Han, 2010), causing it to be tagged as low-priority information under incidental encoding conditions. Second, the basal ganglia–prefrontal gating circuit itself is functionally compromised, as evidenced by imbalanced amygdala-cortical connectivity, a preponderance of limbic input with relatively deficient executive control, and abnormal alpha-band inhibitory processes. Recent causal evidence further indicates that the transition from attention to working memory relies on active inhibitory control mechanisms (Liu et al., 2025); in subthreshold depression, a negative self-concept may drive an excessive engagement of this inhibitory gating on self-face identity. Consequently, in the surprise trial, where no explicit memory expectation exists, the combination of a weakened salience signal and excessive gating inhibition prevents self-face identity from being encoded into working memory. Critically, once an explicit expectation is introduced in control trials, encoding recovers rapidly, confirming that the deficit is specific to incidental encoding conditions and that the basic capacity for encoding remains intact.

### Implications for Understanding Self-Referential Processing in Subthreshold Depression

The present findings reveal a critical dissociation in subthreshold depression: early attentional capture (Lee et al., 2024; Jublie & Kumar, 2021) and explicit recognition of the self-face (Quevedo et al., 2018) remain intact, yet incidental encoding of self-face identity is selectively compromised. This localizes the deficit to the gating stage between perception and working memory consolidation (cf. Firestone & Scholl, 2015) and extends Beck’s (1976) cognitive model by demonstrating that negative self-concept disrupts even the pre-conscious prioritization of self-relevant information. Notably, this deficit cannot be attributed to the confound of emotional valence: by employing neutral self-faces, the present study circumvents the blunted emotional reactivity characteristic of depression (McIvor et al., 2021; Rottenberg et al., 2005) and reveals that altered self-processing in subthreshold depression is fundamental and valence-general, consistent with evidence from depressed adolescents (Alarcón et al., 2019). Finally, the rapid recovery of encoding in control trials indicates that this “de-privileging” of the self-face is malleable, raising the possibility that interventions enhancing the intrinsic salience of self-related cues could mitigate progression to major depressive disorder.

### Limitation

Notably, in Experiment 2, location report accuracy on the surprise trial was modestly reduced relative to pre-surprise and control trials, likely because the location judgment was administered after the surprise question (i.e., identity), causing some participants to forget the location while responding to the surprise question (Chen & Wyble, 2016). This reduction was absent in Experiment 1, perhaps owing to the lower cognitive demands of discriminating simple animal drawings compared with faces.

One potential concern is that identity report accuracy for other-face trials did not immediately recover to ceiling on the first control trial in Experiment 2. It is important to note, however, that accuracy on the remaining control trials (Control 2–4) was consistently high for both self-face and other-face conditions, confirming that participants were fully capable of encoding face identity once the expectation to report had been established (Chen & Wyble, 2015a, 2016). The modest performance on Control Trial 1 may reflect the additional cognitive demands of reconfiguring encoding strategies for complex face stimuli immediately following the unexpected surprise probe (Chen et al., 2019). Critically, the surprise trial remains the key index for assessing selective encoding failure under incidental conditions, as it captures memory performance in the absence of any prior expectation to report the probed attribute. The selective drop in self-face accuracy on the surprise trial, which was disproportionate relative to other-face accuracy in the subthreshold depression group, is therefore not compromised by the transient recovery delay on the first control trial.

## Conclusion

In conclusion, using the attribute amnesia paradigm, the present study systematically investigated the incidental encoding of self-face identity in individuals with subthreshold depression. Results revealed a selective enhancement of the attribute amnesia effect for self-face identity, whereas no such deficit emerged for nonsocial stimuli or for other-faces, indicating that the impairment is specific to self-relevant information. Mechanistically, this encoding failure appears to arise from a negative self-concept that weakens the intrinsic salience of the self-face, a deficit that is further compounded by functional abnormalities in the basal ganglia–prefrontal gating circuit. This gating process relies on active inhibitory control mechanisms; in subthreshold depression, these mechanisms may become excessively engaged for self-relevant information. Critically, the rapid recovery of encoding in control trials indicates that the basic capacity for encoding remains intact. The present study thus pinpoints the self-processing deficit in subthreshold depression to the gating stage between attention and consolidation, thereby providing novel empirical support for extending cognitive models of depression to the earliest, automatic levels of self-relevant processing.

## Authors’ contributions

Writing: Mao Wen and Binyuan Su. Programming: Mao Wen, Binyuan Su and Tianyi Gu. Data collection: Mao Wen, Binyuan Su and Yufan Chen. Analysis: Mao Wen and Binyuan Su. Review and editing: Pengmin Qin. Funding acquisition: Pengmin Qin. All authors approved the final version of the manuscript for submission.

## Funding

This research was supported by the National Key Research and Development Program of China (2025YFE0213500), the National Science Foundation of China (32371098), Guangdong Basic and Applied Basic Research Foundation (2024A1515011429) and the Research Center for Brain Cognition and Human Development, Guangdong, China (No. 2024B0303390003).

## Data availability

Data or materials for the experiments are available from the corresponding author upon request.

## Code availability

All code generated or used during the study are available from the corresponding author upon reasonable request.

## Declarations

### Conflict of interest

The authors have no relevant financial or non-financial interests to disclose.

### Ethics approval

This study was performed in line with the principles of the Declaration of Helsinki. Approval was granted by the Ethics Committee of the School of Psychology, South China Normal University (No. SCNU-PSY-2025-353).

### Consent to participate

Informed consent was obtained from all individual participants included in the study.

### Consent for publication

The authors affirm that human research participants provided informed consent for publication of the images in Figure 2b.

## Notes

### Competing Interest Statement

The authors have declared no competing interest.

